# Characterizing the effect of aging on resting and event-evoked ocular response dynamics

**DOI:** 10.64898/2026.05.06.723160

**Authors:** Mert Huviyetli, Claudia Contadini-Wright, Maria Chait

## Abstract

Ocular measures are increasingly used as non-invasive proxies of cognitive processes such as attention and listening effort. However, their interpretation in aging populations is complicated by concurrent changes in ocular physiology and oculomotor control, raising a critical question: to what extent do age-related differences in these measures reflect cognitive rather than other physiological factors? Here, we dissociate these contributions by characterizing ocular dynamics (resting and event-evoked) during passive fixation in younger (N = 98, 18–35 years) and older adults (N = 71, 60+ years). Aging is associated with pronounced alterations in pupil dynamics, including reduced baseline variability and slower, attenuated responses to both auditory and visual events. In contrast, microsaccade dynamics did not correlate with aging. Across measures, ocular responses showed moderate-to-high within-subject stability across blocks, and factor analysis in the older cohort revealed separable components reflecting instantaneous pupil responsivity, sustained pupil responsivity, and microsaccade dynamics, with additional variance associated with sensory decline and age-related changes in pupil dynamics. Together, these findings demonstrate a clear dissociation: pupil-based metrics are strongly influenced by aging, whereas microsaccades remain comparatively stable across age groups. This dissociation provides a principled basis for interpreting ocular indices in aging research and highlights the need to account for baseline physiological differences when inferring cognitive processes from eye-based measures.

## Introduction

Ocular dynamics provide a unique, non-invasive window into cognitive processes and have attracted increased attention in hearing science in recent years as measures of listening effort and attention allocation (Zekveld et al., 2011; Widmann et al., 2014; Van der Wel and Van Steenbergen, 2018; Zhao et al., 2019c; Magliacano et al., 2020; Schneider et al., 2021; Chen et al., 2023; Contadini-Wright et al., 2023; Coupal et al., 2025; LaCroix and Ratiu, 2025; Huviyetli and Chait, 2026; Liu and Chait, 2026).

The most widely studied ocular measure is pupil dilation because of its long-established link to locus coeruleus (LC) activity: baseline pupil diameter (PD) is thought to reflect tonic LC activity associated with general alertness, whereas transient dilations index phasic activations that signal rapid arousal in response to novel stimuli (Aston-Jones and Cohen, 2005; Wang and Munoz, 2015; Joshi and Gold, 2020).

Recently, microsaccades (MS) have also attracted attention as potentially complementary measures of perceptual engagement: MS are small, involuntary fixational eye movements that are thought to support automatic exploration of the visual environment (Martinez-Conde et al., 2006; Otero-Millan et al., 2008). Novel stimuli transiently suppress MS, a phenomenon known as microsaccadic inhibition (MSI) (Engbert and Kliegl, 2003; Rolfs et al., 2008). This suppression is modulated by stimulus salience and attentional allocation (White and Rolfs, 2016; Roberts et al., 2019; Zhao et al., 2019c). MSI is considered to reflect the operation of a rapid, adaptive orienting mechanism that interrupts ongoing processing to prioritise novel inputs. The generation of MS relies on a distributed network involving the superior colliculus (SC), frontal eye fields (FEF), and visual cortex (Munoz and Istvan, 1998; Hafed et al., 2009; Hafed et al., 2015). The same network is associated with the control of larger saccades; We specifically focus on MS here because of its fixational nature and the fact that it can be measured alongside PD.

Growing evidence indicates that PD and MS reflect at least partially dissociable processes (Contadini-Wright et al., 2023; Zhao et al., 2024; Huviyetli and Chait, 2026; Liu and Chait, 2026). For example, MS effects tend to be confined to temporal windows in which attention is actively engaged (Liu and Chait, 2026), such as the reduction in MS rate observed during the presentation of behaviourally relevant keywords (Contadini-Wright et al., 2023). By contrast, PD responses are often more sustained, consistent with longer-lasting arousal-related activity. Taken together, these findings suggest that PD and MS provide complementary indices of the engagement of attentional and arousal systems during task performance. Relative to methods such as EEG, ocular measures are also attractive from a practical perspective because they are easier and less costly to acquire, e.g. in clinical settings. In addition, multiple indices can be measured simultaneously during fixation, including PD, MS, and blinks, making eye tracking a particularly efficient tool for assessing cognitive performance.

An important line of inquiry, particularly relevant in the context of hearing research, concerns age-related changes in cognitive and sensory processing (Piquado et al., 2010; Zekveld et al., 2011; Zhao et al., 2019b; He et al., 2020; Herrmann and Ryan, 2024). However, aging affects both the structure of the eyes and their neural control (Salvi et al., 2006; Kovács, 2022), which may complicate the interpretation of eye tracking results.

To assess these concerns, here we examined and compared simple ocular responsivity (pupil dilation, blinks, and microsaccades) measured during fixation in standard laboratory settings in younger (<35 years old) and older (>60 years old) participants. Quantifying such baseline differences is essential: if an experimental manipulation is taken to reflect differences in cognitive processing between younger and older groups, it is necessary to demonstrate that the observed effects exceed, and cannot be reduced to, basic age-related changes in ocular physiology or oculomotor control.

Two opposing branches of the autonomic nervous system (ANS) regulate pupil size: the parasympathetic pathway, which mediates constriction, and the sympathetic pathway, which governs dilation (McDougal and Gamlin, 2015). Pupillometric measures such as the light reflex and resting pupil size are widely used to index the functional integrity of these autonomic pathways. With aging, pupil dynamics undergo marked alterations, including reduced resting pupil size (senile miosis), slower constriction and dilation responses, and attenuated light reflex amplitude (Bitsios et al., 1996b; Muppidi et al., 2013; Wang et al., 2016; Tekin et al., 2018). These age-related changes arise from multiple, interacting sources. Structural alterations in the iris musculature and reduced efficiency of autonomic innervation impair the mechanical and neural control of pupil responses (Salvi et al., 2006; Kovács, 2022). In parallel, neuromodulatory systems that are closely linked to pupil diameter—most notably the locus coeruleus–norepinephrine (LC-NE) and basal forebrain– acetylcholine systems—are particularly vulnerable to age-related neurodegeneration (Aston-Jones and Cohen, 2005; Mather and Harley, 2016; Larsen and Waters, 2018; Joshi and Gold, 2020; Bañuelos et al., 2023). Because these neuromodulatory systems regulate global arousal and vigilance—states that are closely indexed by pupil diameter—their age-related decline is expected to contribute to altered pupil dynamics during aging.

To our knowledge, evidence on how aging affects MS is limited. Port et al. (2016) examined MS across the lifespan (from age 4 to 66) during complex visual search tasks. They found that while visual search performance (i.e., the speed of finding a target) follows a U-shaped function—peaking around age 20 and declining in both younger children and older group—the rate of MS generation shows only a very slight increase across the lifespan. This suggests that the fundamental oculomotor system responsible for MS is preserved in aging, even as the cognitive and sensory processes involved in visual search efficiency fluctuate with age. Oh et al. (2026) examined MS frequency across distinct age cohorts (20s, 60s, and 70s) during passive fixation and reported a significant age-related increase in MS frequency. Crucially, within the older groups (60s and 70s), individuals with lower cognitive performance, *as measured by the Mini-Mental State Examination*, exhibited significantly higher MS rates compared to cognitively unimpaired peers of the same age group.

To support the interpretation of age-related findings, and to determine whether age-related changes in PD are associated with changes in MS dynamics, we provide a detailed characterisation of ocular responses, including PD and MS, measured during rest and in response to brief visual and auditory stimuli. Specifically, we aimed to: (1) provide large-sample distributional data for basic ocular measures; (2) characterize the relationships among basic ocular indices and their associations with age and hearing loss; and (3) examine the correspondence between visually evoked and auditory evoked ocular responses.

## Materials and Methods

### Ethics

The research was approved by the Research Ethics Committee of University College London. Participants provided written informed consent and were paid for their participation.

### Participants

We collected data from 98 young adults aged between 18–35 years (mean age = 23.32 years, 70 females) and 71 older adults over 60 years old (mean age = 69.54 years, 40 females) (Figure 1). The dataset was compiled from five separate experiments (three studies with younger adults and two with older adults), where an “Oculometrics” measurement block was included prior to the main experiment. This oculometrics block forms the basis of the present study. All participants reported no history of major surgery within the past two years and no known neurological disorders. Hearing profiles were assessed in the older participants; younger participants self-reported normal hearing and no history of hearing problems.

**Figure 1.**
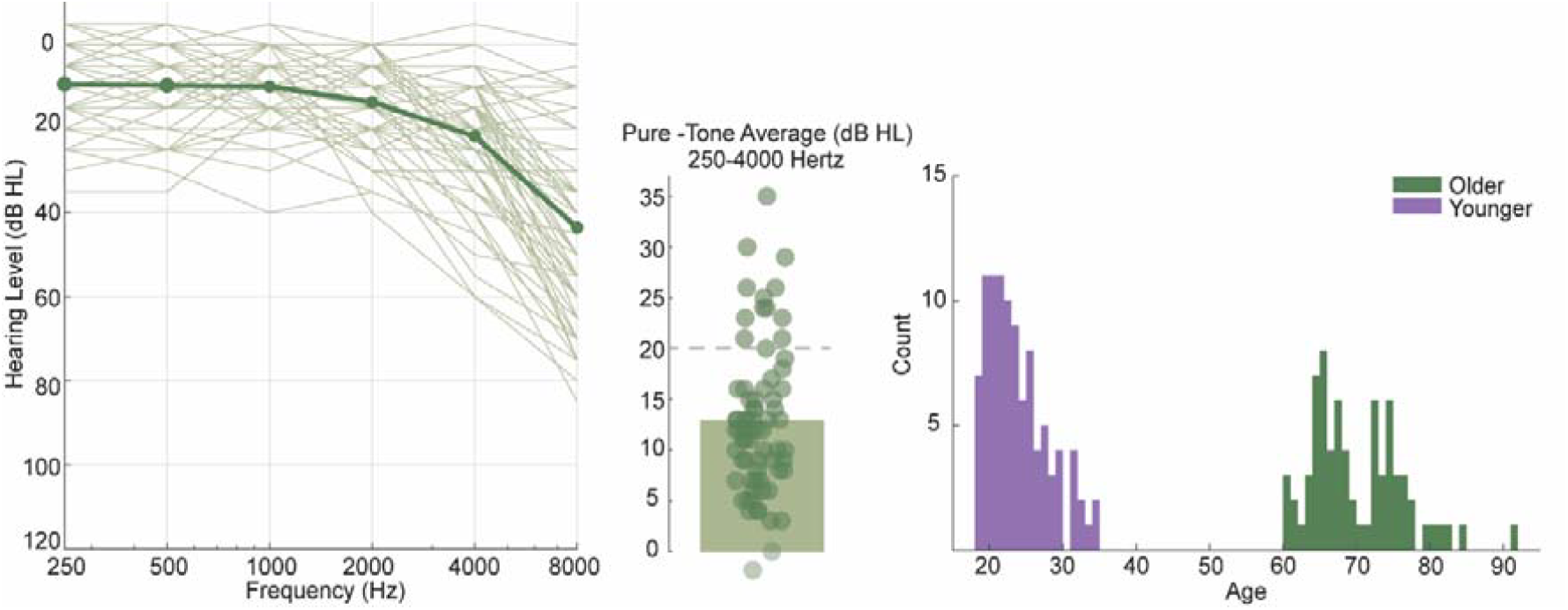
Age and hearing-threshold characteristics of the participants. Left: Individual audiograms (older participants) showing hearing level (dB HL) between 250–8000 Hz. Thin lines represent individual participants, and the thick line denotes the group average. Middle: Distribution of pure-tone average (PTA; 250–4000 Hz), with individual participants shown as dots and the bar indicating the mean PTA across listeners. The dashed line represents the upper limit of clinically normal hearing (20 dB HL). Right: participant age distribution.

Older participants did not report hearing-aid use and were selected to include individuals with no worse than mild hearing loss, defined as a pure-tone average (PTA) threshold of 40 dB HL or less across frequencies from 250 Hz to 4 kHz in the better ear. Audiometric assessments were conducted for 0.25, 0.5, 1, 2, 4, and 8 kHz. In one study (n = 39), hearing thresholds were measured with the GSI Pello audiometer using Radioear DD45 supra-aural headphones. In the other study (n = 32), thresholds were assessed with the Interacoustics AC-629 audiometer paired with Sennheiser HDA300 headphones. As expected, older participants showed elevated hearing thresholds at higher frequencies (Slade et al., 2020). Most participants had a PTA of 20 dB HL or lower (see Figure 1, middle), which falls within the *‘clinically defined normal hearing range*.*’*

### Procedure

The experimental session lasted approximately 15 minutes and included five ocular measures. Throughout the sessions, the default screen background was grey (RGB: 127.50, 127.50, 127.50), see the luminance value below.

#### Resting pupil diameter

Absolute pupil size (mm) was recorded while participants fixated on a black cross presented at the centre of a grey background screen (RGB: 127.50, 127.50, 127.50) during a 30-second trial. Resting pupil diameter was defined as the mean pupil size over this period. Pupil size variability was quantified as the standard deviation within the same period. Normalized variability (Coefficient of Variation) was calculated by dividing the variability value by the corresponding resting mean pupil size.

Note that resting pupil data were not available for one of the young cohorts (n = 32). Therefore, analyses involving resting pupil measures in the younger group were conducted on a subsample of 66 participants.

#### Gradient screen test

This task involved a gradual change in screen brightness across five levels: white (255, 255, 255), light grey (191.25, 191.25, 191.25), grey (127.50, 127.50, 127.50), dark grey (63.75, 63.75, 63.75), and black (0, 0, 0). Values indicate RGB color codes. Luminance was measured from the chinrest position using a Konica Minolta LS-150 luminance meter. The measured luminance levels were 114.2 cd/m^2^ for white, 63.34 cd/m^2^ for light grey, 26.66 cd/m^2^ for grey, 5.40 cd/m^2^ for dark grey, and 0.59 cd/m^2^ for black. The entire sequence lasted approximately 50 seconds, with each color presented for 10 seconds. The pupil response for each luminance condition was calculated as the mean absolute pupil size (mm) during the final 5 seconds of each trial. Pupil diameter range was determined by subtracting the mean pupil diameter during the black condition from that during the white condition.

#### Flash-evoked ocular responses

Ocular response to a sudden white or black flash were quantified. Each trial began with a grey screen serving as baseline. A brief 300 ms flash—either white or black—was then presented, after which the screen returned to grey. White and black flashes were presented in separate blocks, each consisting of 30 trials and lasting approximately 3 minutes.

#### Sound-evoked pupil response

Pupil responses to 500-ms harmonic tones (fundamental frequency = 200 Hz, 30 harmonics) were recorded to investigate auditory-evoked ocular dynamics. Each block consisted of 30 trials and lasted approximately 3.5 minutes.

The gradient screen test was always presented first, while the remaining conditions were randomized. Participants sat with their head fixed on a chinrest in front of a monitor (24-inch BENQ XL2420T with a resolution of 1920×1080 pixels and a refresh rate of 60 Hz) in a dimly lit and acoustically shielded room (IAC triple walled sound-attenuating booth). They were instructed to continuously fixate on a black cross presented at the centre of the screen. An infrared eye-tracking camera (Eyelink 1000 Desktop Mount, SR Research Ltd.) placed below the monitor at a horizontal distance of 62cm from the participant was used to record eye data. The standard five-point calibration procedure for the Eyelink system was conducted prior to each measure, and participants were instructed to avoid any head movement after calibration. During the experiment, the eye-tracker continuously tracked gaze position and recorded pupil diameter, focusing binocularly at a sampling rate of 1000 Hz. Participants were instructed to blink naturally. Prior to each trial, the eye tracker automatically checked that the participants’ eyes were open and fixated appropriately; trials would not start unless this was confirmed.

Auditory stimuli were presented diotically via a Roland Tri-Capture 24-bit/96 kHz soundcard connected to Sennheiser HD558 headphones. The loudness of the stimuli was individually adjusted to ensure a comfortable listening level for each participant.

### Pupillometry Preprocessing and Analysis

Where possible, data from the left eye were analysed. Intervals during which the participant gazed away from fixation (i.e., outside a radius of 100 pixels around the centre of the fixation cross), as well as periods of full or partial eye closure (e.g., blinks), were automatically treated as missing data. For blink-related artefacts, a window extending 150 ms before and 150 ms after blink onset/offset was removed and subsequently reconstructed using shape-preserving piecewise cubic interpolation. Epochs with more than 50% missing data or those determined to be particularly noisy were removed from the data. For each participant, the pupil response was time-domain averaged across all epochs to produce a single time series per condition.

### Pupil Rate Analysis

To derive the pupil dilation rate (PDR) and constriction rate (PCR) time series, pupil events were extracted from the continuous, smoothed data (150-ms Hanning window). Based on Joshi et al. (2016) and Zhao et al. (2019c; 2024), the events were defined as local minima that are followed by continuous dilation/constriction of the pupil for at least 100 ms. In each condition, for each participant, the event time series was summed and normalized by the number of trials and the sampling rate. Then, a causal smoothing kernel ω⍰(τ⍰)=α⍰^2×⍰τ⍰×⍰e^(-ατ) was applied with a decay parameter of α⍰=⍰1/50 ms (Dayan and Abbott, 2005; Rolfs et al., 2008; Widmann et al., 2014) paralleling a similar technique for computing neural firing rates from neuronal spike trains (Dayan & Abbott, 2005; see also Joshi et al., 2016; Rolfs et al., 2008). The mean across trials was computed and baseline corrected. To account for the time delay caused by the smoothing kernel, the time axis was shifted by the latency of the peak of the kernel window.

### Microsaccade Preprocessing and Analysis

Intervals during which full or partial eye closure was detected (e.g., during blinks) were automatically treated as missing data and were not interpolated. For microsaccade (MS) analyses, data within 100 ms before and after blink onset and offset were excluded to avoid blink-related artefacts. A shorter exclusion window was used in the MS analysis because pupil data exhibit slow dynamics, whereas microsaccades are transient events.

MS detection was based on an approach proposed by Engbert and Kliegl (2003). MS were extracted from the continuous eye-movement data based on the following criteria: (1) A velocity threshold of λ = 6 times the median-based standard deviation within each condition (2) Above-threshold velocity lasting between 5ms and 100 ms (3) The events are detected in both eyes with onset disparity <10 ms (4) The interval between successive microsaccades is longer than 50ms. Extracted microsaccade events were represented as unit pulses (Dirac delta). For each participant, the event time series was summed and normalized by the number of trials and the sampling rate. The microsaccade rate was then computed in the same way as described for the pupil dilation rate, above.

### Blink Analysis

Sharp decreases in the pupil size signal, including both full and partial eye closures, were classified as blinks. Partial eye closure events were operationally defined as instances in which pupil size fell below ±7 SD relative to the mean. For each detected blink, a ±100 ms window (i.e., 100 ms before and after the identified time point, similar to MS analysis) was included to capture the full blink-related signal change. Blink occurrences were then identified on a trial-by-trial basis and averaged at each time point, yielding a time-resolved blink incidence measure (percentage of trials containing a blink at each time point) for subsequent analyses.

## Statistical analysis

In addition to analyzing absolute pupil size (in mm) and Microsaccade rate (in Hz), these measures were also normalized for each participant and condition. This approach accounts for individual and age-related differences in baseline ocular activity, allowing analyses to focus on relative changes and dynamic response patterns.

For PD data, normalization involved calculating the mean and standard deviation of the relevant signal during the 1-second baseline period (prior to flash/sound onset) across all trials and using these values to z-score the data. For MS data, for each participant and condition, was first smoothed (see details above), and z-scoring was then applied using the mean and standard deviation computed over the 1-second baseline period.

To identify time intervals showing significant differences between age groups, we employed a non-parametric, bootstrap-based between-group analysis (Efron and Tibshirani, 1994). For each bootstrap iteration (1,000 iterations with replacement), participants were resampled independently within each group, and the group-averaged difference time series (Younger − Older) was computed. This procedure yielded a distribution of mean group differences at each time point. Time points were considered statistically significant if more than 95% of the bootstrap samples lay consistently above or below zero. This analysis was performed across the entire epoch.

Normality of the data was assessed using the Kolmogorov–Smirnov test. Where assumptions of normality were met, independent-samples t-tests were used for two-group comparisons; otherwise, Mann–Whitney U tests were applied. These tests were conducted on condition- or time-window–averaged ocular metrics.

For the older group, an exploratory factor analysis was performed to investigate the underlying structure of the ocular measures, age, and hearing loss variables (see more details below). Factors were extracted using principal components analysis with Varimax rotation and Kaiser normalization. Based on the Kaiser criterion of eigenvalues greater than 1, five factors were retained. The rotated solution converged in five iterations, and factor loadings smaller than 0.30 were suppressed in the factor matrix to aid interpretation.

Follow-up Spearman’s rank-order correlations were calculated within the older group to evaluate the relationships between ocular measures and age, and the associations between ocular measures and hearing thresholds.

### The parameters used in these analyses were

1. Age
2. Hearing acuity (pure-tone average; PTA in the worse ear): Quantified such that higher values signify better hearing.
3. Mean pupil size (during resting state)
4. Pupil size range (black–white screen)
5. Variability in pupil size (during resting state)
6. Sound evoked mean PD: 0-2.25 s (relative to sound onset)
7. Sound evoked PD peak: 0-2.25 s (relative to sound onset)
8. (8) Sound evoked mean MS rate: 0-0.5 s (relative to sound onset)
9. Mean MS rate (during baseline): −1-0 s (1 s pre-sound onset)
10. Constriction onset time (defined as the latency of a sustained negative slope (< −0.30 mm/s for ≥ 5 ms)
11. Maximum constriction velocity (MCV), defined as the peak negative slope;
12. Maximum constriction (mm)
13. T75, defined as the time from stimulus onset to 75% recovery of the pupil response (relative to baseline);
14. DV1, the re-dilation velocity at 1 s after the minimum pupil response (mean slope within ±50 ms);
15. Late re-dilation velocity, defined as the peak positive slope occurring after the T75 timepoint.

For measures of pupil instantaneous dynamics (parameters 10-15) we focused on the white flash-evoked reflex. The various parameters are illustrated in Figure 4B. All timing measures are reported relative to stimulus onset and absolute pupil size is used (mm).

## Results

### Resting-state pupil dynamics change with aging, whereas microsaccades and blinks remain unaltered

Figure 2 (top plots) shows the mean absolute pupil size, variance and normalized variability over a 30-second resting-state period. Older participants exhibited significantly smaller (t (131) = 3.718, p < .001) and less variable absolute pupil size compared to younger participants (U = 965.000, Z = −5.607, p < .001, variability and U = 1153.000, Z = −4.761, p < .001, normalized variability).

**Figure 2.**
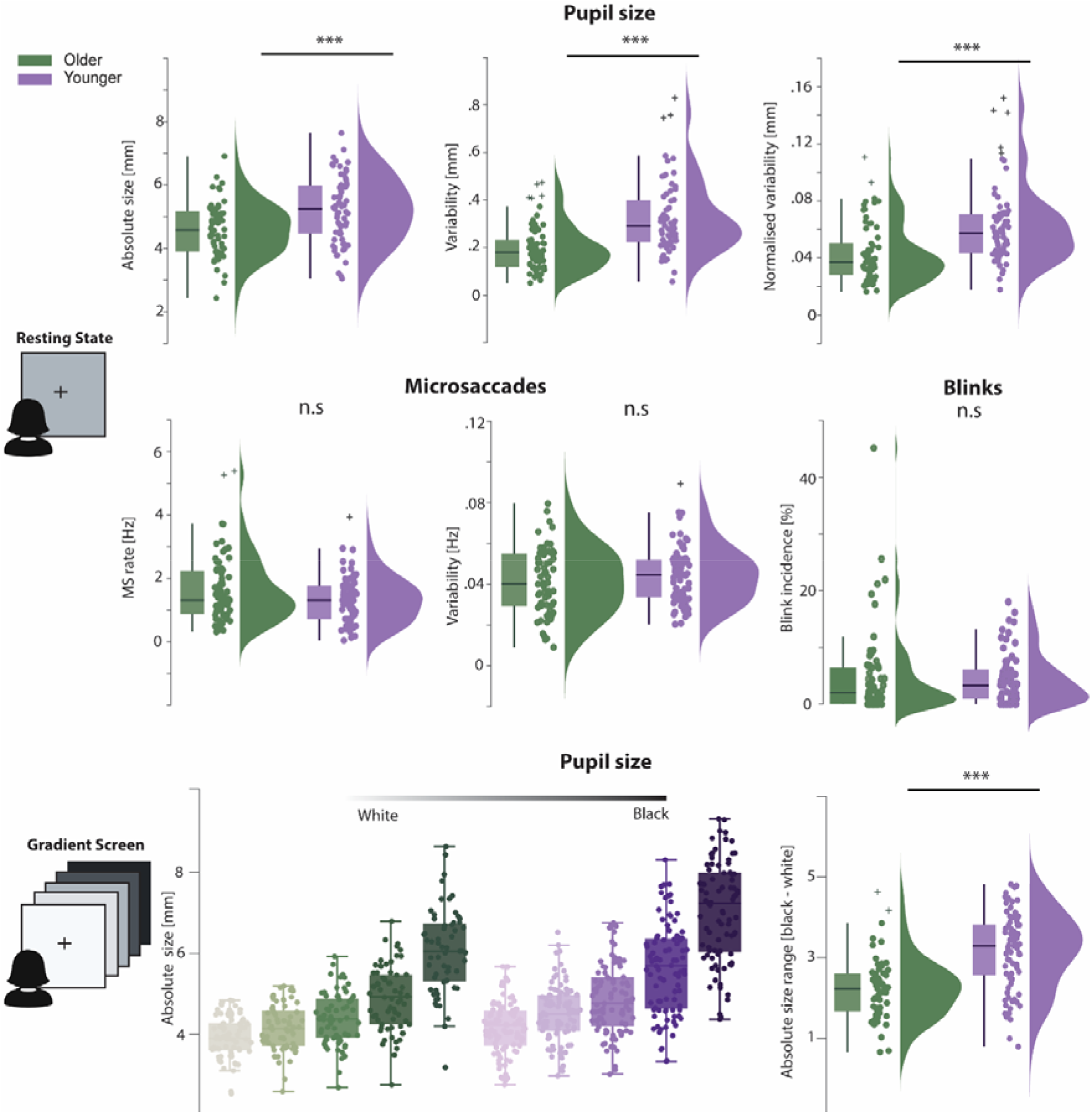
Resting state pupil, microsaccade and blink dynamics: **Top:** Group distributions of mean absolute pupil size, within-subject variability, and within-subject normalized variability. Dots represent individual participants; boxplots show median and interquartile range. **Middle:** Group distributions of mean Microsaccade rate, Microsaccade rate within-subject variability, and blink incidence. **Bottom left:** Absolute pupil size as a function of screen brightness, showing larger pupil size for darker backgrounds and overall larger pupils in the younger cohort. Bottom Right: Absolute pupil range (black–white difference). *** indicates p < .001

The middle panels of Figure 2 show MS and blink measures during the resting-state period. Resting MS rate did not differ between age groups (U = 1945.50, Z = −1.33, p = .183), though between-subject variability was larger in the older group. Within-individual MS variance was also not different (t(132) = −1.24, p = .216). Similarly, the incidence of blinks (%) did not differ between groups (U = 2101.00, Z = −0.64, p = .521).

Figure 2 (bottom plots) illustrates changes in pupil size as a function of screen brightness, manipulated across five levels (see methods). A Linear Mixed-Effects Model (LMM) was conducted to examine the effects of Brightness (five levels: white, light grey, grey, dark grey, black) and Age (two groups: Older, Younger) on pupil size, while accounting for within-subject correlations using a random intercept for participants. The model included main effects of Brightness and Age, as well as their two-way interaction. The LMM revealed a statistically significant main effect of Brightness on pupil size, F(4, 637.237) = 575.219, p < .001, indicating that mean pupil size differed significantly across the five luminance conditions. A significant main effect of Age was also observed, F(1, 168.145) = 24.799, p < .001, suggesting overall differences in pupil diameter between the Older and Younger groups. Additionally, a significant two-way interaction between screen brightness and age group was observed, F(4, 637.24) = 14.24, p < .001. This interaction indicates that the effect of screen brightness on pupil size differed significantly between younger and older groups. As illustrated in Figure 2, both age groups showed an increase in pupil size from white to black screens. However, this increase was markedly steeper in the younger group. This divergence resulted in a widening gap between age groups, with younger participants exhibiting a significantly larger pupil size than older participants in the black screen condition. To better quantify this, pupil size range was computed by subtracting the value for the white screen condition from that of the black screen. Older participants exhibited a significantly smaller pupil size range compared to younger participants (t(140) = 5.537, p < .001), confirming a reduced dynamic pupil response to luminance changes in the older group.

Overall, these findings align with previous literature reporting age-related decline in baseline pupil size, variability, and range. Such differences may reflect age-associated changes in the autonomic nervous system, which regulates pupil size, and/or structural alterations such as age-related atrophy of the pupil muscle (Pfeifer et al., 1983; Winn et al., 1994; Manaye et al., 1995; Bitsios et al., 1996a; Mather and Harley, 2016; Tekin et al., 2018; Telek et al., 2018; Zhao et al., 2019a). In contrast, MS and blink incidence did not differ systematically between groups, although the older group contained more outlier participants with higher rates (see discussed below).

### Reduced variability and slower pupil responses to sudden events (auditory and visual) in the older group

Figure 3 (top row) illustrates pupil responses to sudden events - *white flashes, black flashes and an auditory burst (Harmonic tone)*. The white flash evoked a rapid constriction, followed by a gradual dilation. A black flash evoked an initial pupil dilation followed by constriction, and then a recovery response. The sound-evoked PD response followed a similar pattern to previous findings (Zhao et al., 2019a), with a rapid increase in pupil size following the sound. In all cases, the younger group exhibited earlier dilation and constriction and a larger overall pupil response (Figure 3, top panels).

**Figure 3.**
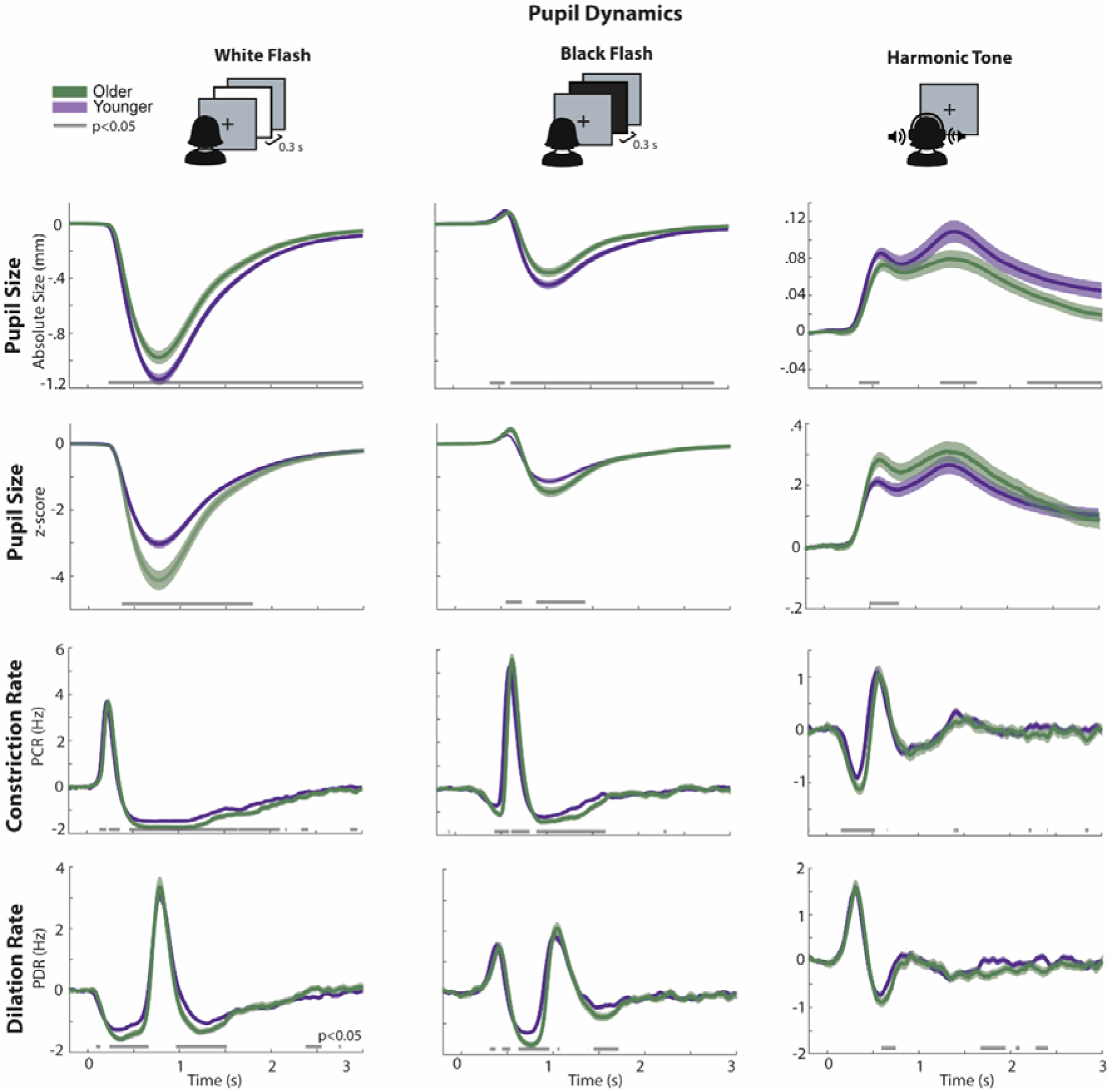
Pupil responsivity to sudden visual and auditory events. **Left panels** show responses to white flashes; **middle panels** show responses to black flashes**; right panels** show responses to the sound. In each set, plots display (1) absolute pupil size, (2) z-scored pupil responses, (3) constriction rate, and (4) dilation rate. Shaded areas indicate SEM. Black bars beneath the traces denote time intervals with significant group differences (p < .05).

**Figure 4.**
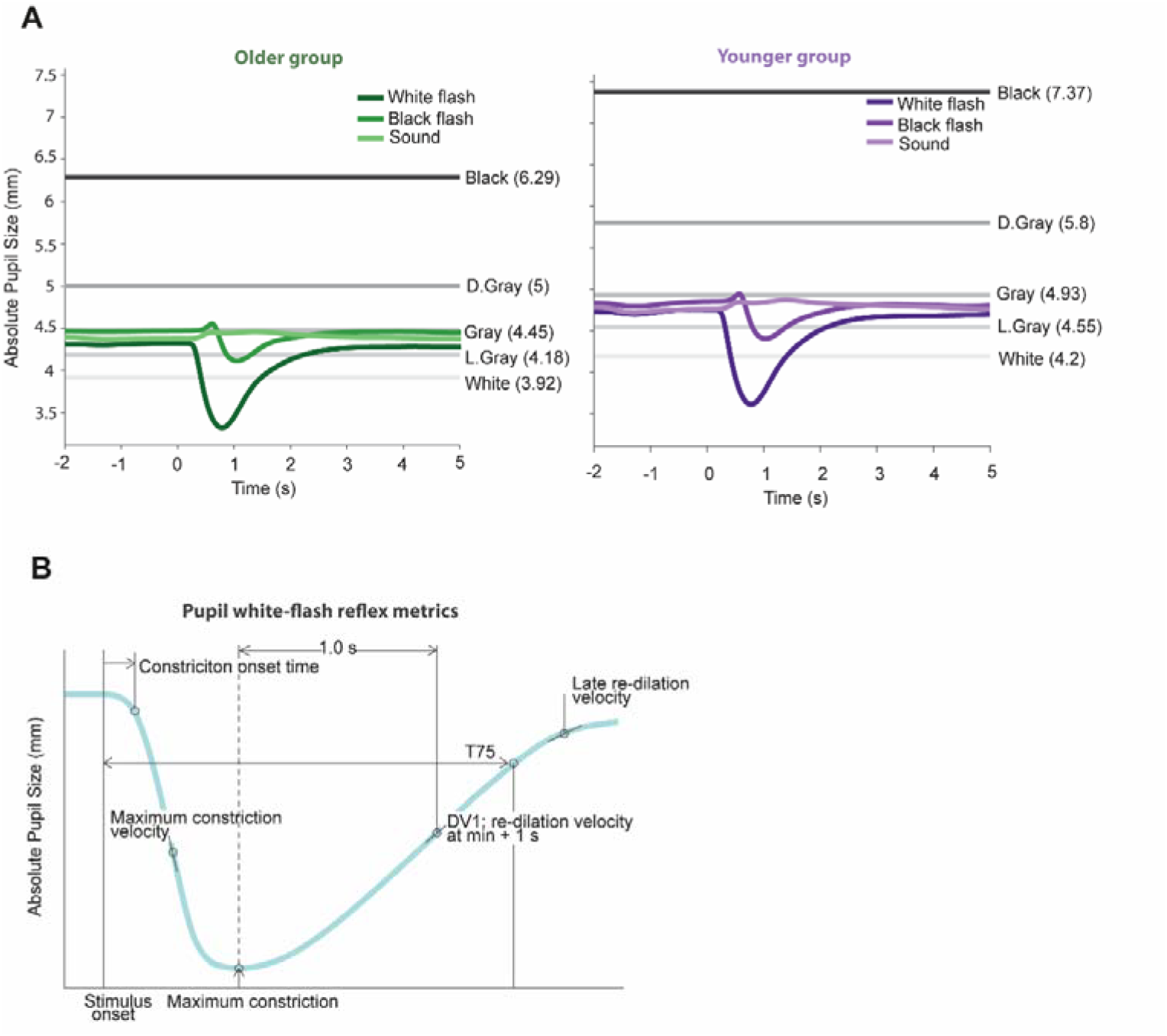
Summary of Pupil responses (in mm) **A:** Time course of mean absolute pupil size (mm) in response to sudden events in the older (left panel, green) and younger (right panel, purple) groups. Horizontal lines indicate pupil sizes measured at different luminance levels (white to black; “screen gradient test”), with corresponding values indicated on the right side of each panel. **B:** A schematic illustration of key metrics derived from the pupil-light reflex (White Flash) to be used for additional analyses below.

These results are consistent with previous findings of age-related changes in pupillary dynamics (Bitsios et al., 1996a; Fotiou et al., 2007; Wang et al., 2016; Tekin et al., 2018; Wang et al., 2018) reporting a reduction in constriction velocity, maximum constriction amplitude, and constriction onset time. These age-related changes are interpreted as arising from a weaker response of the iris muscles in the older group, potentially due to age-related changes in autonomic nervous system function and/or iris muscle elasticity.

Further, we converted the data to z-scores using the within-subject mean and standard deviation during the baseline period (1 second before flash/sound onset). This normalisation allowed us to interpret the results relative to each participant’s own baseline pupil dynamics. The z-normalized data show a generally opposite pattern (between groups) to that observed when expressing the signals in mm (see: Figure 3 z-score data). This is because, similar to what was seen for the resting state (Figure 2), older participants show lower baseline variability compared to the younger group. Mann–Whitney U tests confirmed that older participants exhibited significantly lower variability during the baseline period across measures (white flash: U = 1691, Z = −5.595, p < .001; black flash: U = 2002, Z = −4.482, p < .001; sound: U = 1965, Z = −4.822, p < .001). Lower variability during baseline means that even small changes in pupil size in mm-scale can translate into relatively large changes in standardized (z-scored) pupil responses.

Beyond overall pupil size changes, direct measures of pupil responsivity, quantified as dilation/constriction rate, may more closely reflect differential engagement of sympathetic (dilation-related, PDR) and parasympathetic (constriction-related, PCR) activity (Joshi et al, 2016; Zhao et al., 2024; Basgol et al., 2025; Huviyetli and Chait, 2026; Liu and Chait, 2026). These measures quantify discrete pupil dilation and constriction events independently of overall changes in pupil size. In all three conditions, we observed faster responsivity (earlier PDR and PCR) in the younger group compared to the older group (Figure 3, bottom).

Figure 4A summarizes the main pupil data for the younger and older groups (in mm), showing the mean pupil-diameter range, as measured during the screen gradient test, and how the phasic responses map onto that range. This should facilitate comparisons with other experiments and may also be useful when assessing specific populations, by indicating how their performance maps onto these group means. Figure 4B illustrates the key measures derived from the White-flash reflex to be used in the Factor and Correlation analyses (below).

### Higher baseline microsaccade rate in older people during flash/sound burst blocks

In contrast to the observation during the resting state block, the older group exhibited a higher baseline MS rate than the younger group (Figure 5). This is consistent with previous findings (Port et al., 2016). Both bootstrap analysis and the Mann–Whitney U test confirmed significant group differences for baseline (−1 to 0 s before events) white-flash (U = 2602, Z = −2.53, p = .011), black-flash (U = 2416, Z = −3.14, p = .002), and HT (U = 2481, Z = −3.18, p = .001). An inspection of the data indicates that the observed difference between the resting-state block and the flash/sound burst blocks is largely attributable to slightly elevated MS baseline activity in the older cohort. Although the underlying cause is uncertain, one possibility is that abrupt sensory events, compared with the resting-state condition, elicit greater active exploration in older participants.

**Figure 5.**
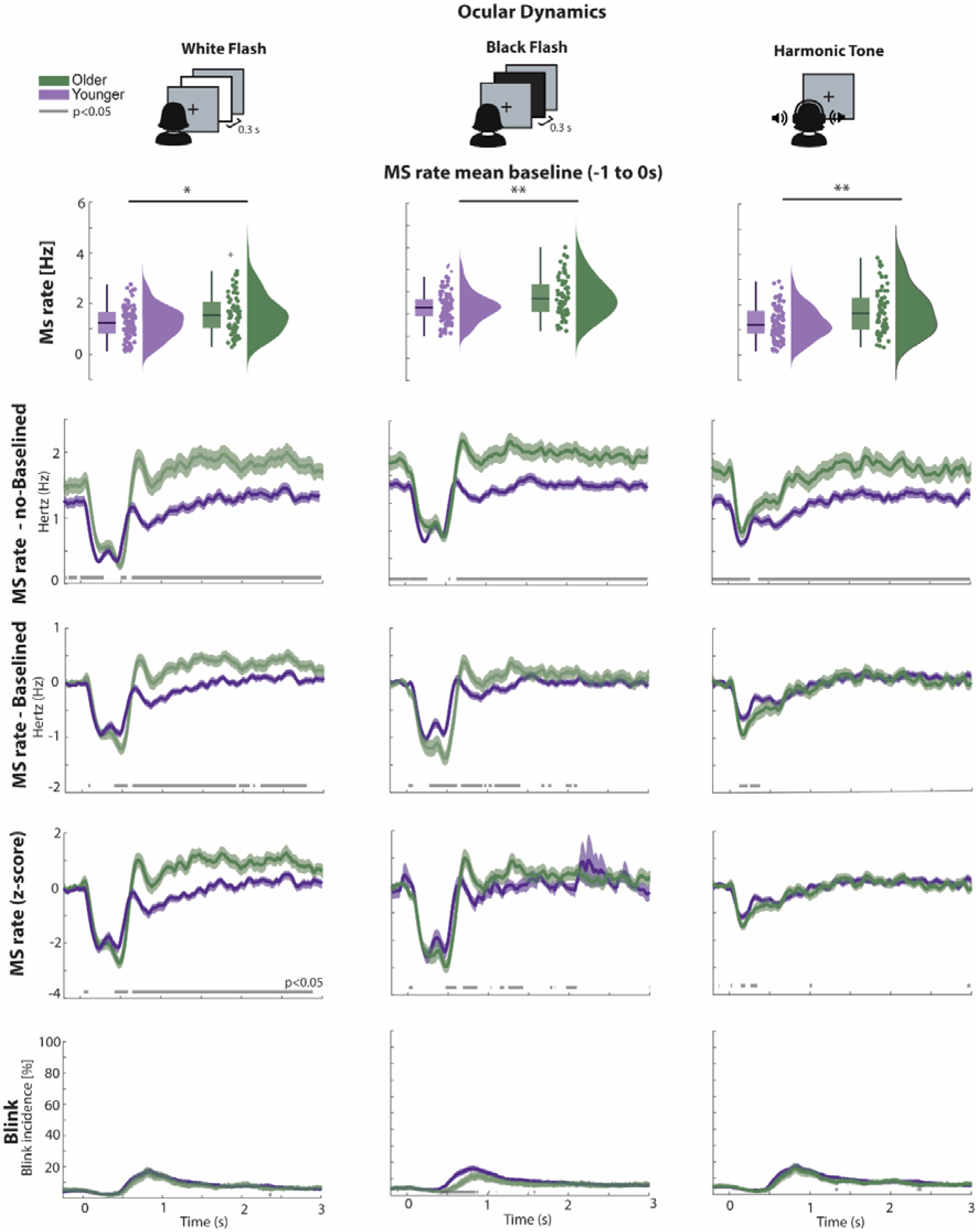
Microsaccade (MS) and Blink responses to sudden events. **Upper panels** indicate baseline mean 1 second before the events. * indicates p < .05; ** p < .01 **Left panels** show responses to white flashes; **middle panels** show responses to black flashes. **right panels** show responses to the sound. In each set, plots display (1) Microsaccade rate (no-baselined (2)Microsaccade rate (baselined) (3)Microsaccade rate (z-scored), and (4) blink (%). Shaded areas indicate SEM.

This baseline difference is crucial for interpreting stimulus-evoked effects. Baseline correction altered the apparent magnitude of MSI (Figure 5, baseline-corrected data). Following correction, MSI appeared more pronounced in the older group; however, this effect is driven mainly by their higher baseline activity rather than by intrinsically stronger MS inhibition.

Indeed, as expected, visual flashes and abrupt auditory bursts elicited robust MSI in younger and older participants (Figure 5;Engbert & Kliegl, 2003; Rolfs et al., 2008). In the flash conditions, the inhibition phase was characterized by two distinct troughs, reflecting the sequential luminance transitions (grey to white/black and back to grey). This was followed by a rebound in MS rate (where activity exceeded that during baseline the baseline rate) before a gradual return to baseline. In contrast, in the HT condition, inhibition was not followed by a rapid rebound; instead, MS rate recovered more gradually over time. A similar pattern was observed in previous work with younger participants (Zhao et al., 2019c; Zhao et al., 2024; Huviyetli and Chait, 2026; Liu and Chait, 2026). This difference in post-inhibitory dynamics evoked by visual and auditory stimuli may reflect modality-specific processing characteristics. In visual processing, the rebound, manifested as an abrupt increase in MS dynamics, may be associated with visual reorienting, whereas similar processes may not be engaged following abrupt auditory events.

Importantly, during the peak inhibition window (MSI trough), MS rates were comparable across groups, suggesting preserved early inhibitory mechanisms in the older group despite their elevated baseline MS activity. However, post-inhibitory recovery was slower in the older group in all three conditions. MSI recovery is often interpreted as reflecting a release of attentional capture (Liu & Chait, 2026; Contadini-Wright et al, 2023), therefore the pattern observed here may be suggesting age-related slowing of attentional re-orienting mechanisms.

Age-related differences in MSI latency were condition-dependent. In the white-flash condition, the older group exhibited earlier MSI onset compared to younger adults (Figure 5). In contrast, in the black-flash condition, this pattern was reversed, with younger group showing earlier inhibition. In the HT (sound) condition, no significant latency differences were observed between age groups.

To further account for baseline differences and standard deviation during the baseline period (similar to the pupil response), we z-transformed the MS data. The z-scored MS results closely mirrored the raw-rate pattern, indicating that the observed age effects were not solely driven by differences in baseline dynamics.

Finally, we analyzed blink responses (Figure 5, bottom). Across all conditions, blink incidence showed a similar overall pattern, characterized by a temporary increase beginning around 500 ms after event onset. Given that event duration was 300 ms for the visual flashes and 500 ms for the sounds, this rise roughly coincides with event offset and is consistent with the previously reported “release of blinking” phenomenon (Fukuda, 1994; Oh et al., 2012; Murali and Händel, 2021). This effect is thought to reflect a core capacity to strategically time blinks away from periods of heightened processing demand. Blink responses to white flash and sound (HT) did not differ significantly between the younger and older group. However, a significant age difference emerged for black flashes. This selective effect suggests that aging may disproportionately affect luminance-decrement (OFF-pathway) processing, whereas luminance-increment (ON-pathway) processing remains relatively preserved.

### Ocular measures show moderate stability across task blocks

We further examined the extent to which baseline mean and variability of pupil-size and microsaccade rate are correlated across blocks (Table 1). Establishing such associations provides an important indication of whether this activity is sufficiently stable within individuals to be considered trait-like, or whether it is primarily state-dependent.

**Table 1.** Spearman’s correlations across sensory conditions in terms of mean and variability of ocular measures. *** indicates p<.001

| mean Baseline Pupil size (-1 to 0 s) |  |  |  |  |
| --- | --- | --- | --- | --- |
|  | White flash | Black flash | Sound | Resting |
| White flash | — |  |  |  |
| Black flash | .764<sup>***</sup> | — |  |  |
| Sound | .861<sup>***</sup> | .795<sup>***</sup> | — |  |
| Resting | .810<sup>***</sup> | .757<sup>***</sup> | .840<sup>***</sup> | — |
| within-subject Pupil size variability (-1 to 0 s) |  |  |  |  |
|  | White flash | Black flash | Sound | Resting |
| White flash | — |  |  |  |
| Black flash | .368<sup>***</sup> | — |  |  |
| Sound | .505<sup>***</sup> | .433<sup>***</sup> | — |  |
| Resting | .488<sup>***</sup> | .541<sup>***</sup> | .516<sup>***</sup> | — |
| mean Microsaccade rate (-1 to 0 s) |  |  |  |  |
|  | White flash | Black flash | Sound | Resting |
| White flash | — |  |  |  |
| Black flash | .748<sup>***</sup> | — |  |  |
| Sound | .763<sup>***</sup> | .766<sup>***</sup> | — |  |
| Resting | .688<sup>***</sup> | .700<sup>***</sup> | .635<sup>***</sup> | — |
| within-subject Microsaccade rate variability (-1 to 0 s) |  |  |  |  |
|  | White flash | Black flash | Sound | Resting |
| White flash | — |  |  |  |
| Black flash | .352<sup>***</sup> | — |  |  |
| Sound | .383<sup>***</sup> | .464<sup>***</sup> | — |  |
| Resting | .241<sup>**</sup> | .366<sup>***</sup> | .338<sup>***</sup> | — |

**Table 2.** Rotated Component Matrix: Extraction method: Principal component analysis. Rotation method: Varimax with Kaiser normalisation. Rotation converged in 5 iterations. Loadings < .30 are suppressed.

| Rotated Component Matrix |  |  |  |  |  |
| --- | --- | --- | --- | --- | --- |
| Variable | Component |  |  |  |  |
|  | Instantaneous pupil responsivity | Sustained pupil responsivity | Aging affected pupil dynamics | Microsaccade dynamics | Sensory decline |
| Age |  |  | **-.780** |  |  |
| Hearing acuity |  |  | **.762** |  | **.336** |
| Resting pupil mean |  | **.725** | **.301** |  |  |
| Pupil size range (black–white screen) |  | **.667** |  |  |  |
| Variability of resting pupil size |  | **.375** | **.510** |  |  |
| Sound-evoked PD peak |  | **.934** |  |  |  |
| Sound-evoked PD mean |  | **.908** |  |  |  |
| Maximum constriction velocity | **.957** |  |  |  |  |
| Constriction onset time | **-.560** |  | **-.535** |  | **.315** |
| Maximum constriction | **-.934** |  |  |  |  |
| T75, recovery time |  |  |  |  | **.918** |
| Late re-dilation velocity | **.713** |  |  |  | **-.474** |
| DV1, re-dilation velocity | **.883** |  |  |  |  |
| Sound-evoked mean MS rate |  |  |  | **.837** |  |
| Mean MS rate (baseline in sound) |  |  |  | **.839** |  |

Across all participants (younger and older groups combined), strong correlations were observed for baseline mean responses in both PD and MS-rate (r = .75–.86, all p < .001). In contrast, within-subject variability showed more moderate correlations (r=.35–.51, all p < .001).

### Comparing auditory and visual evoked MSI

Figure 6 compares MSI responses across the sound and the two flash conditions. Although the salience of auditory and visual events is difficult to compare, MSI in both age groups, elicited by sounds and flashes, showed broadly similar temporal profiles. Notably, an earlier MSI in response to sound, compared to flashes, was seen (this is particularly apparent in the younger group), consistent with previous findings (Rolfs 2008).

**Figure 6.**
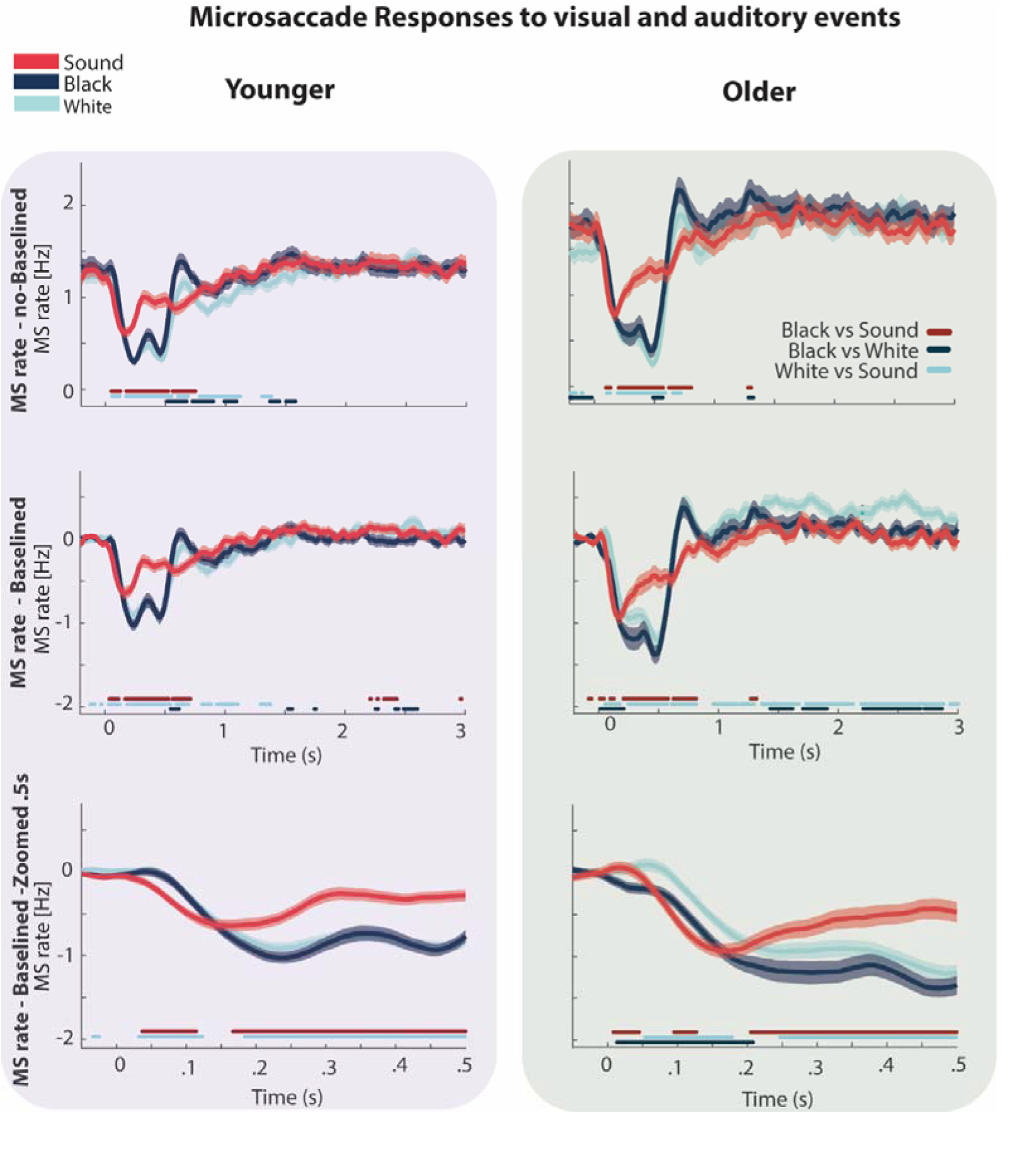
Microsaccade responses across stimulus modalities in younger and older group. **Left panels (purple shaded)** show responses for younger adults; **right panels (green shaded)** show responses for older group. Top row shows non-baselined MS rate; middle row shows baseline-corrected MS rate; bottom row shows a zoomed view of the baseline-corrected MS rate within the first 0.5 s post-stimulus. Time courses are shown with shaded areas indicating SEM. Coloured bars at the bottom of each panel denote time intervals with significant pairwise differences between conditions (p< .05; see legend below the traces in top right panel)

### Dissociable components of aging, pupil dynamics, and microsaccade measures in older adults

Focusing on the older group only (N = 71), a principal component analysis (PCA) was conducted to examine how the aging, hearing, and ocular metrics grouped together (see methods). In line with previous work using light-reflex paradigms as markers of autonomic nervous system activity (Wang et al., 2016; Tekin et al., 2018), we focused on white flash– evoked pupil response features (see Figure 4B), alongside resting pupil measures and sound-evoked PD and MS responses (see detailed in Methods). Overall the analysis included the following parameters: **age, hearing acuity** (PTA in the worse ear), resting pupil metrics (mean size, range, variability), **sound-evoked responses** (mean and peak pupil dilation, mean MS rate, and baseline MS rate), and **white flash–evoked pupil dynamics** (see Figure 4B and methods). Baseline-corrected data were used for all ocular measures, except for those pertaining to baseline dynamics. Pupil size data are in mm.

The KMO measure was 0.65, and Bartlett’s test of sphericity was significant (p < .001), indicating that the data are suitable for component analysis. Rotation converged after five iterations and yielded a five-component solution. Together, the extracted components explained 77.6% of the total variance in the data. Components 1 to 5 accounted for 23.8%, 19.8%, 13.1%, 10.8%, and 9.9% of the variance, respectively. The rotated component matrix is shown in Table 3.

**Table 3.** Spearman Correlations Between Age and Other (audiometric and ocular) Measures, r = Spearman rank correlation coefficient. ANS = autonomic nervous system; MS = microsaccade; PTA = pure-tone average e* p < .05 ** p < .01. *** p < .001.

| Variable | r | p | Justification |
| --- | --- | --- | --- |
| Resting pupil mean | -.482*** | .000041 | Effect of aging on pupil responsivity |
| Pupil size range (black–white screen) | -.231 | .078334 | Effect of aging on pupil responsivity |
| Variability of resting pupil size | -.406*** | .000723 | Effect of aging on pupil responsivity |
| Normalised variability of resting pupil size | -.288* | .019048 | Effect of aging on pupil responsivity |
| Hearing acuity (Worse Ear, PTA) | -.526*** | .000002 | Effect of aging on hearing thresholds |
| Sound-evoked PD peak | -.319*** | .006788 | Effect of aging on tone-evoked pupil response |
| Sound-evoked mean PD | -.319*** | .006702 | Effect of aging on tone-evoked pupil response |
| Sound-evoked mean MS rate | .058 | .631480 | Effect of aging on tone-evoked MS response |
| Mean MS rate (baseline in sound) | .093 | .442891 | Effect of aging on ocular dynamics |
| Maximum constriction velocity | -.264* | .027452 | Effect of aging on ANS activity |
| Constriction onset time | .368** | .001749 | Effect of aging on ANS activity |
| Maximum constriction | .219 | .069148 | Effect of aging on ANS activity |
| T75, recovery time | .163 | .177837 | Effect of aging on ANS activity |
| Late re-dilation velocity | -.283* | .017654 | Effect of aging on ANS activity |
| DV1, re-dilation velocity | -.270* | .023640 | Effect of aging on ANS activity |

The first component captured aspects of instantaneous pupil responsivity, including measures such as onset time and maximum velocity. The second component reflected sustained pupil size and pupil-size variability, with strong loadings from resting pupil diameter as well as sound-evoked baseline and peak pupil measures.

The third component was associated with age-related factors, with the strongest loadings from age and hearing thresholds. This component also loaded on baseline pupil variability and response time. Notably, it appeared to be linked more closely to variability measures than to mean pupil size, suggesting that age may be more strongly associated with fluctuations in pupil dynamics than with pupil size itself.

The fourth component was defined by microsaccade-related measures, indicating that microsaccade dynamics represent an independent dimension that is largely dissociable from both age-related and pupil-related metrics.

The fifth component was most strongly characterised by the redilation T75 variable, reflecting the speed of the pupil response, together with other latency-related measures. There was also some indication of an association with hearing status, despite these measures not having been acquired during auditory presentation.

### Aging is associated with reduced pupil size and responsivity

As a follow-up to the analyses above, we conducted correlation analyses to examine the relationships between age (older group; >60 years), hearing sensitivity, and ocular metrics.

Table 1 presents the Spearman correlations between age and all study variables. Age was significantly associated with multiple variables, including several measures of pupil responsivity, consistent with a large body of work reporting changes to pupil responsivity with aging (Bitsios et al., 1996a; Tekin et al., 2018; Telek et al., 2018; Zhao et al., 2019b). Notably, however, age was not significantly correlated with any of the MS measures.

No correlations (corrected for age) were observed between hearing status and any of the measures.

In summary, the conclusion from this rather large dataset (N=71) of older adults aged 60-92 confirms robust links between aging and pupil dynamics but no correlation with hearing loss. It is also noteworthy that baseline MS incidence did not correlate with age, despite the group-level differences observed. This suggests that although baseline MS rates differ broadly between ages 18–30 and 60+, they do not appear to track aging closely in the same way that PD does.

## Discussion

Ocular dynamics are increasingly used to probe cognitive processes and are likely to play an even greater role in future cognitive and neuroscience research (He et al., 2020; Schneider et al., 2021; Contadini-Wright et al., 2023; Herrmann et al., 2025; Liu and Chait, 2026). However, interpreting these measures presents an important challenge, as aging can influence ocular responses independently of cognitive processing. Consequently, it is often unclear whether observed differences reflect changes in cognitive mechanisms or age-related alterations in basic sensory or oculomotor responses. In the present study, we aimed to address this issue by characterizing ocular indices measured during fixation to simple auditory and visual events, and by examining how these responses—and their relationship across modalities—vary with aging. Consistent with previous reports, our results showed that pupil responses (including sustained size and various measures of instantaneous responsivity) changed with aging. In contrast, MS dynamics did not correlate with aging.

### Pupil dynamics change with ageing

Pupil size reflects the dynamic balance between two opposing branches of the autonomic nervous system: the parasympathetic pathway, which drives constriction, and the sympathetic pathway, which mediates dilation (McDougal and Gamlin, 2015). Accordingly, basic pupil measures such as resting pupil size and the light reflex provide non-invasive indices of autonomic and neuromodulatory function (Aston-Jones and Cohen, 2005; Larsen and Waters, 2018; Joshi and Gold, 2020). Aging is associated with pronounced alterations in pupil dynamics—including smaller resting pupil size, slower constriction and dilation, and reduced reflex amplitude—likely reflecting both peripheral changes in the iris and age-related alterations in neuromodulatory systems (Bitsios et al., 1996a; Mather and Harley, 2016; Tekin et al., 2018; Wang et al., 2018; Zhao et al., 2019b). Consistent with this, our resting pupil measures showed that the older group exhibited a smaller resting pupil size and a narrower pupil dynamic range. In addition, pupil responses to sudden events (white– black flashes and sound) revealed reduced velocity, amplitude of both pupil constriction and dilation, and delayed onset of the pupil response in the older group. These changes likely reflect weaker iris muscle responses with aging, potentially arising from alterations in autonomic nervous system function and/or reduced elasticity of the iris musculature.

A particularly noteworthy observation is that of reduced baseline pupil size variability observed in the older group. Neural variability, *once considered noise*, is now understood as a fundamental property of brain function that supports cognitive flexibility and adaptive behaviour, with higher variability linked to better cognitive performance (Grady and Garrett, 2018). Neural variability is thought to be influenced by neuromodulatory systems such as dopamine and noradrenaline, as well as by the balance between excitation and inhibition in neural circuits (Waschke et al., 2017), both of which change with aging. The Locus coeruleus (LC)–noradrenaline system is particularly relevant here, as it regulates cortical gain and moment-to-moment neural fluctuations, and is closely associated with pupil dynamics (Aston-Jones and Cohen, 2005; Joshi and Gold, 2020). With age, the LC shows reduced structural integrity and diminished noradrenergic output, and these changes are associated with decline in attention, working memory, and cognitive flexibility (Mather and Harley, 2016; Dahl et al., 2019). The reduced baseline pupil variability observed in older adults may therefore reflect, in part, diminished LC-driven flexibility in arousal.

From a practical perspective, reduced pupil variability influences comparisons between younger and older groups when responses are expressed as z-score-normalized values. Because z-scoring entails division by variance, the lower variance observed in the older group can make their responses appear larger overall. Increased phasic pupil responses in the older group demonstrate that, relative to their own baseline activity, older adults exhibit greater pupil responsivity to abrupt events such as flashes or auditory stimuli, an important consideration when interpreting group differences. Therefore, presenting both z-scored responses and raw responses in millimetres is useful for obtaining a more complete perspective on responsivity.

### Microsaccade dynamics show a mixed effect with ageing

Unlike PD, literature on the effects of aging on MS dynamics is scarce. Port et al. (2016) showed that MS rate increases only slightly across the lifespan during a complex visual search task, indicating that the fundamental oculomotor mechanisms underlying MS generation remain relatively intact with age. Recently, Oh et al. (2026) reported a significant age-related increase in MS frequency during passive fixation, and further demonstrated that individuals with lower mini-mental test scores exhibited significantly higher MS rates within the same age group.

We quantified MS rate during resting state and during passive exposure to sounds and flashes. We found that resting-state MS rate and variability did not differ between younger and older groups, suggesting that basic MS dynamics are largely preserved with aging. However, in the event-evoked blocks (white–black flashes and sound), older participants showed a higher baseline MS rate compared to younger participants. These findings suggest that while the basic microsaccade-generating system is preserved with aging, stimulus-driven contexts may reveal age related differences. Future work will be needed to clarify the sources of this variability and the mechanisms shaping individual MS dynamics.

In response to transient visual and auditory stimuli, we observed a robust MSI in both groups. Overall, MSI responses to auditory and visual events followed a broadly similar temporal pattern in younger and older group. However, the magnitude of this inhibition appeared larger in the older group, largely due to their higher baseline MS rate.

### PCA and correlation analyses within the older group confirm a strong link between pupil dynamics and aging, whilst MS dynamics appear independent

The factor analysis of the measures examined here yielded a five-component solution that explained a substantial proportion of the variance in the older group (>77%), indicating that the dataset captures several robust and dissociable dimensions of performance and physiology.

The first two components mapped specifically onto instantaneous and sustained pupil responsivity, providing strong evidence that these aspects of the pupil response reflect separable mechanisms. This distinction is important because it suggests that studies relying on pupillometry should not treat pupil responsivity as a unitary construct.

The third component, associated with aging, clustered with measures of sustained pupil activity, including pupil size and variability, as well as constriction response time. The association with pupil variability is especially compelling, both in light of the discussion above concerning z-scoring and because it aligns with previous reports of age-related changes in broader neural variability (Waschke et al., 2017; Grady and Garrett, 2018). Taken together, these findings suggest that variability is a meaningful dimension of age-related physiological change. Practically, this argues strongly for incorporating pupil size variability measures into experiments with older listeners, so as to fully characterize their responsivity.

Component 4 was defined by MS-related measures and was clearly independent of all other components, demonstrating that MS dynamics are distinct from both pupil dynamics and aging. This independence is theoretically and methodologically important. It indicates that MS dynamics are not confounded by age-related changes in ocular physiology and therefore may offer a particularly stable basis for comparing cognitive functioning between younger and older listeners.

Perhaps most intriguingly, the final factor, which we term “sensory decline,” grouped hearing-loss measures with indices of pupil responsivity to light flashes, including constriction time and multiple measures reflecting the speed of return to baseline. Notably, this association emerged only in the dimension-reduction analysis and was not evident in the partial correlations, suggesting that the shared variance between hearing loss and pupil responsivity becomes apparent only once the broader covariance structure among measures is taken into account. This pattern nonetheless points to a specific relationship between hearing loss and pupil responsivity, with practical implications for the design and interpretation of studies seeking to relate pupillometry to hearing loss. The effects were relatively modest in the present sample, likely because most participants had only mild hearing loss, and should be re-examined in samples with a broader range of auditory impairment.

## Conclusions

This study provides distributional data from a relatively large sample of both older and younger participants. These data should be valuable for researchers working with smaller samples, as they offer a normative framework against which individual participants or cohorts can be compared.

Overall, and in line with previous work, we observed widespread age-related differences across pupil-dilation measures, including responses to sound, as well as correlations with age even within the older cohort itself. These findings reinforce the conclusion that pupil dynamics are strongly shaped by aging. This has important implications for interpretation: when pupil-based measures are used to index perceptual challenge or cognitive effort across age groups, age-related physiological differences may substantially confound those inferences.

By contrast, MS dynamics, which are attracting increasing interest as measures complementary to pupil dilation, were not correlated with pupil-dilation measures or with aging. This independence suggests that MS dynamics may provide a useful index of cognitive processing without the same concern about age-related physiological constraints. In that sense, they may offer a particularly valuable tool for cross-age comparisons.

Interestingly, some measures of pupil responsivity that were not sound-evoked were associated with hearing loss. Although this effect requires further investigation, it raises the possibility that certain aspects of basic pupil responsivity may be sensitive to sensory decline in ways that are separable from task-evoked responses.

An important caveat is that we interpret this brief measurement series as reflecting relatively automatic pupil responsivity, because participants were not required to make any overt response to the stimuli. However, this assumption should be treated with some caution. Older participants may have been more motivated or more engaged than younger participants (Ryan and Campbell, 2021), potentially reflecting greater compliance, stronger interest in the study, or a greater tendency to follow experimental instructions carefully. Even in a paradigm involving only passive viewing and listening, differences in engagement or vigilance could plausibly influence ocular measures, particularly pupil responses, through changes in arousal or attentional state (Hopstaken et al., 2015). Thus, although the passive nature of the task minimizes explicit performance demands, it does not fully eliminate the possibility that age-group differences in motivation or engagement contributed to the observed effects.

## Acknowledgements

MH is supported by a PhD studentship from the Ministry of National Education of the Republic of Turkiye. CCW and MC are supported by the NIHR UCLH BRC Deafness and Hearing Problems Theme. The funders had no role in study design, data collection, and analysis, the decision to publish, or preparation of the manuscript. We thank Miaohsu Chang for her contribution to data collection from the older participant group.

## References

Aston-Jones, G., and Cohen, J.D. (2005). An integrative theory of locus coeruleus-norepinephrine function: adaptive gain and optimal performance. Annu Rev Neurosci 28, 403–450. doi: 10.1146/annurev.neuro.28.061604.135709.

Bañuelos, C., Kittleson, J.R., LaNasa, K.H., Galiano, C.S., Roth, S.M., Perez, E.J., et al. (2023). Cognitive Aging and the Primate Basal Forebrain Revisited: Disproportionate GABAergic Vulnerability Revealed. J Neurosci 43(49), 8425–8441. doi: 10.1523/jneurosci.0456-23.2023.

Basgol, H., Dayan, P., and Franz, V.H. (2025). Violation of auditory regularities is reflected in pupil dynamics. Cortex 183, 66–86. doi: 10.1016/j.cortex.2024.10.023.

Bitsios, P., Prettyman, R., and Szabadi, E. (1996a). Changes in autonomic function with age: a study of pupillary kinetics in healthy young and old people. Age Ageing 25(6), 432– 438. doi: 10.1093/ageing/25.6.432.

Bitsios, P., Szabadi, E., and Bradshaw, C. (1996b). The inhibition of the pupillary light reflex by the threat of an electric shock: a potential laboratory model of human anxiety. Journal of psychopharmacology 10(4), 279–287.

Chen, S., Epps, J., and Paas, F. (2023). Pupillometric and blink measures of diverse task loads: Implications for working memory models. British Journal of Educational Psychology 93, 318–338.

Contadini-Wright, C., Magami, K., Mehta, N., and Chait, M. (2023). Pupil dilation and microsaccades provide complementary insights into the dynamics of arousal and instantaneous attention during effortful listening. Journal of Neuroscience 43(26), 4856–4866.

Coupal, P., Zhang, Y., and Deroche, M. (2025). Reduced Eye Blinking During Sentence Listening Reflects Increased Cognitive Load in Challenging Auditory Conditions. Trends Hear 29, 23312165251371118. doi: 10.1177/23312165251371118.

Dahl, M.J., Mather, M., Düzel, S., Bodammer, N.C., Lindenberger, U., Kühn, S., et al. (2019). Rostral locus coeruleus integrity is associated with better memory performance in older adults. Nat Hum Behav 3(11), 1203–1214. doi: 10.1038/s41562-019-0715-2.

Dalmaso, M., Castelli, L., Scatturin, P., and Galfano, G. (2017). Working memory load modulates microsaccadic rate. Journal of Vision 17(3), 6–6. doi: 10.1167/17.3.6.

Dayan, P., and Abbott, L.F. (2005). Theoretical neuroscience: computational and mathematical modeling of neural systems. MIT press.

Efron, B., and Tibshirani, R.J. (1994). An introduction to the bootstrap. CRC press.

Engbert, R., and Kliegl, R. (2003). Microsaccades uncover the orientation of covert attention. Vision research 43(9), 1035–1045.

Fotiou, D.F., Brozou, C.G., Tsiptsios, D.J., Fotiou, A., Kabitsi, A., Nakou, M., et al. (2007). Effect of age on pupillary light reflex: evaluation of pupil mobility for clinical practice and research. Electromyogr Clin Neurophysiol 47(1), 11–22.

Fukuda, K. (1994). Analysis of eyeblink activity during discriminative tasks. Percept Mot Skills 79(3 Pt 2), 1599–1608. doi: 10.2466/pms.1994.79.3f.1599.

Grady, C.L., and Garrett, D.D. (2018). Brain signal variability is modulated as a function of internal and external demand in younger and older adults. Neuroimage 169, 510–523. doi: 10.1016/j.neuroimage.2017.12.031.

Hafed, Z.M., Chen, C.-Y., and Tian, X. (2015). Vision, perception, and attention through the lens of microsaccades: mechanisms and implications. Frontiers in systems neuroscience 9, 167.

Hafed, Z.M., Goffart, L., and Krauzlis, R.J. (2009). A neural mechanism for microsaccade generation in the primate superior colliculus. science 323(5916), 940–943.

He, M., Heindel, W.C., Nassar, M.R., Siefert, E.M., and Festa, E.K. (2020). Age-related changes in the functional integrity of the phasic alerting system: a pupillometric investigation. Neurobiology of Aging 91, 136–147. doi: 10.1016/j.neurobiolaging.2020.02.025.

Herrmann, B., and Ryan, J.D. (2024). Pupil size and eye movements differently index effort in both younger and older adults. Journal of Cognitive Neuroscience 36(7), 1325– 1340.

Herrmann, B., Scharf, F., and Widmann, A. (2025). Eye movements of younger and older adults decrease during story listening in background noise. Hear Res 468, 109447. doi: 10.1016/j.heares.2025.109447.

Hopstaken, J.F., van der Linden, D., Bakker, A.B., and Kompier, M.A.J. (2015). The window of my eyes: Task disengagement and mental fatigue covary with pupil dynamics. Biological Psychology 110, 100–106. doi: 10.1016/j.biopsycho.2015.06.013.

Huviyetli, M., and Chait, M. (2026). The Interplay of Bottom-Up Arousal and Attentional Capture during Auditory Scene Analysis: Evidence from Ocular Dynamics. Journal of Neuroscience.

Joshi, S., and Gold, J.I. (2020). Pupil Size as a Window on Neural Substrates of Cognition. Trends Cogn Sci 24(6), 466–480. doi: 10.1016/j.tics.2020.03.005.

Joshi, S., Li, Y., Kalwani, R.M., and Gold, J.I. (2016). Relationships between Pupil Diameter and Neuronal Activity in the Locus Coeruleus, Colliculi, and Cingulate Cortex. Neuron 89(1), 221–234. doi: 10.1016/j.neuron.2015.11.028.

Kovács, I. (2022). Effects of ageing on the eyes. Developments in Health Sciences 4(1), 21– 25.

LaCroix, A.N., and Ratiu, I. (2025). Saccades and Blinks Index Cognitive Demand during Auditory Noncanonical Sentence Comprehension. Journal of Cognitive Neuroscience 37(6), 1147–1172.

Larsen, R.S., and Waters, J. (2018). Neuromodulatory Correlates of Pupil Dilation. Front Neural Circuits 12, 21. doi: 10.3389/fncir.2018.00021.

Liu, X., and Chait, M. (2026). Dissociable Pupil and oculomotor markers of attention allocation and distractor suppression during listening. Journal of Neuroscience 46(1).

Magliacano, A., Fiorenza, S., Estraneo, A., and Trojano, L. (2020). Eye blink rate increases as a function of cognitive load during an auditory oddball paradigm. Neuroscience Letters 736, 135293.

Manaye, K.F., McIntire, D.D., Mann, D.M., and German, D.C. (1995). Locus coeruleus cell loss in the aging human brain: a non-random process. J Comp Neurol 358(1), 79–87. doi: 10.1002/cne.903580105.

Martinez-Conde, S., Macknik, S.L., Troncoso, X.G., and Dyar, T.A. (2006). Microsaccades counteract visual fading during fixation. Neuron 49(2), 297–305.

Mather, M., and Harley, C.W. (2016). The Locus Coeruleus: Essential for Maintaining Cognitive Function and the Aging Brain. Trends Cogn Sci 20(3), 214–226. doi: 10.1016/j.tics.2016.01.001.

McDougal, D.H., and Gamlin, P.D. (2015). Autonomic control of the eye. Compr Physiol 5(1), 439–473. doi: 10.1002/cphy.c140014.

Munoz, D.P., and Istvan, P.J. (1998). Lateral inhibitory interactions in the intermediate layers of the monkey superior colliculus. J Neurophysiol 79(3), 1193–1209. doi: 10.1152/jn.1998.79.3.1193.

Muppidi, S., Adams-Huet, B., Tajzoy, E., Scribner, M., Blazek, P., Spaeth, E.B., et al. (2013). Dynamic pupillometry as an autonomic testing tool. Clinical Autonomic Research 23(6), 297–303.

Murali, S., and Händel, B. (2021). The latency of spontaneous eye blinks marks relevant visual and auditory information processing. J Vis 21(6), 7. doi: 10.1167/jov.21.6.7.

Oh, J., Jeong, S.Y., and Jeong, J. (2012). The timing and temporal patterns of eye blinking are dynamically modulated by attention. Hum Mov Sci 31(6), 1353–1365. doi: 10.1016/j.humov.2012.06.003.

Oh, S., Nairuz, T., Park, S.-J., and Lee, J.-H. (2026). Simultaneous Analysis of Microsaccades and Pupil Size Variations in Age-Related Cognitive Impairment Using Eye-Tracking Technology. Journal of Eye Movement Research 19(2), 29.

Otero-Millan, J., Troncoso, X.G., Macknik, S.L., Serrano-Pedraza, I., and Martinez-Conde, S. (2008). Saccades and microsaccades during visual fixation, exploration, and search: foundations for a common saccadic generator. Journal of vision 8(14), 21–21.

Pfeifer, M.A., Weinberg, C.R., Cook, D., Best, J.D., Reenan, A., and Halter, J.B. (1983). Differential changes of autonomic nervous system function with age in man. Am J Med 75(2), 249–258. doi: 10.1016/0002-9343(83)91201-9.

Piquado, T., Isaacowitz, D., and Wingfield, A. (2010). Pupillometry as a measure of cognitive effort in younger and older adults. Psychophysiology 47(3), 560–569.

Port, N.L., Trimberger, J., Hitzeman, S., Redick, B., and Beckerman, S. (2016). Micro and regular saccades across the lifespan during a visual search of “Where’s Waldo” puzzles. Vision Res 118, 144–157. doi: 10.1016/j.visres.2015.05.013.

Roberts, M.J., Lange, G., Van Der Veen, T., Lowet, E., and De Weerd, P. (2019). The Attentional Blink is Related to the Microsaccade Rate Signature. Cerebral Cortex 29(12), 5190–5203. doi: 10.1093/cercor/bhz058.

Rolfs, M., Kliegl, R., and Engbert, R. (2008). Toward a model of microsaccade generation: The case of microsaccadic inhibition. Journal of vision 8(11), 5–5.

Ryan, A.D., and Campbell, K.L. (2021). The ironic effect of older adults’ increased task motivation: Implications for neurocognitive aging. Psychonomic Bulletin & Review 28(6), 1743–1754. doi: 10.3758/s13423-021-01963-4.

Salvi, S., Akhtar, S., and Currie, Z. (2006). Ageing changes in the eye: this article is part of a series on ageing edited by Professor Chris Bulpitt. Postgraduate medical journal 82(971), 581–587.

Schneider, A., Sonderegger, A., Krueger, E., Meteier, Q., Luethold, P., and Chavaillaz, A. (2021). The interplay between task difficulty and microsaccade rate: Evidence for the critical role of visual load. Journal of Eye Movement Research 13(5), 10.16910/jemr.16913.16915. 16916.

Slade, K., Plack, C.J., and Nuttall, H.E. (2020). The Effects of Age-Related Hearing Loss on the Brain and Cognitive Function. Trends in Neurosciences 43(10), 810–821. doi: 10.1016/j.tins.2020.07.005.

Tekin, K., Sekeroglu, M.A., Kiziltoprak, H., Doguizi, S., Inanc, M., and Yilmazbas, P. (2018). Static and dynamic pupillometry data of healthy individuals. Clinical and Experimental Optometry 101(5), 659–665.

Telek, H.H., Erdol, H., and Turk, A. (2018). The effects of age on pupil diameter at different light amplitudes. Beyoglu Eye J 3(2), 80–85.

Van der Wel, P., and Van Steenbergen, H. (2018). Pupil dilation as an index of effort in cognitive control tasks: A review. Psychonomic bulletin & review 25(6), 2005–2015.

Wang, C.A., and Munoz, D.P. (2015). A circuit for pupil orienting responses: implications for cognitive modulation of pupil size. Curr Opin Neurobiol 33, 134–140. doi: 10.1016/j.conb.2015.03.018.

Wang, C.A., Tworzyanski, L., Huang, J., and Munoz, D.P. (2018). Response anisocoria in the pupillary light and darkness reflex. Eur J Neurosci 48(11), 3379–3388. doi: 10.1111/ejn.14195.

Wang, Y., Zekveld, A.A., Naylor, G., Ohlenforst, B., Jansma, E.P., Lorens, A., et al. (2016). Parasympathetic nervous system dysfunction, as identified by pupil light reflex, and its possible connection to hearing impairment. PloS one 11(4), e0153566.

Waschke, L., Wöstmann, M., and Obleser, J. (2017). States and traits of neural irregularity in the age-varying human brain. Scientific Reports 7(1), 17381. doi: 10.1038/s41598-017-17766-4.

White, A.L., and Rolfs, M. (2016). Oculomotor inhibition covaries with conscious detection. Journal of Neurophysiology 116(3), 1507–1521.

Widmann, A., Engbert, R., and Schröger, E. (2014). Microsaccadic responses indicate fast categorization of sounds: a novel approach to study auditory cognition. J Neurosci 34(33), 11152–11158. doi: 10.1523/jneurosci.1568-14.2014.

Winn, B., Whitaker, D., Elliott, D.B., and Phillips, N.J. (1994). Factors affecting light-adapted pupil size in normal human subjects. Invest Ophthalmol Vis Sci 35(3), 1132– 1137.

Yamagishi, S., and Furukawa, S. (2025). Microsaccade Direction Reveals the Variation in Auditory Selective Attention Processes. The Journal of Neuroscience 45(45), e1623242025. doi: 10.1523/jneurosci.1623-24.2025.

Zekveld, A.A., Kramer, S.E., and Festen, J.M. (2011). Cognitive load during speech perception in noise: the influence of age, hearing loss, and cognition on the pupil response. Ear Hear 32(4), 498–510. doi: 10.1097/AUD.0b013e31820512bb.

Zhao, S., Bury, G., Milne, A., and Chait, M. (2019a). Pupillometry as an Objective Measure of Sustained Attention in Young and Older Listeners. Trends Hear 23, 2331216519887815. doi: 10.1177/2331216519887815.

Zhao, S., Bury, G., Milne, A., and Chait, M. (2019b). Pupillometry as an objective measure of sustained attention in young and older listeners. Trends in hearing 23, 2331216519887815.

Zhao, S., Contadini-Wright, C., and Chait, M. (2024). Cross-modal interactions between auditory attention and oculomotor control. Journal of Neuroscience 44(11).

Zhao, S., Yum, N.W., Benjamin, L., Benhamou, E., Yoneya, M., Furukawa, S., et al. (2019c). Rapid ocular responses are modulated by bottom-up-driven auditory salience. Journal of Neuroscience 39(39), 7703–7714.

